# Untangling the Relatedness among Correlations, Part III: Inter-Subject Correlation Analysis through Bayesian Multilevel Modeling for Naturalistic Scanning

**DOI:** 10.1101/655738

**Authors:** Gang Chen, Paul A. Taylor, Xianggui Qu, Peter J. Molfese, Peter A. Bandettini, Robert W. Cox, Emily S. Finn

**Affiliations:** Scientific and Statistical Computing Core, National Institute of Mental Health, USA; Department of Mathematics and Statistics, Oakland University, USA; Section on Functional Imaging Methods, National Institute of Mental Health, USA

## Abstract

While inter-subject correlation (ISC) analysis is a powerful tool for naturalistic scanning data, drawing appropriate statistical inferences is difficult due to the daunting task of accounting for the intricate relatedness in data structure as well as the intricacy of handling the multiple testing issue. In our previous work we proposed nonparametric approaches to performing group ISC analysis through bootstrapping for one group of subjects and permutation testing for two groups (Chen et al., 2016). A more flexible methodology is to parametrically build a linear mixed-effects (LME) model that captures the relatedness embedded in the data (Chen et al., 2017); in addition, the LME approach also has the capability of incorporating explanatory variables such as subject-grouping factors and quantitative covariates. However, the whole-brain LME modeling methodology still faces some challenges. When an LME model becomes sophisticated, it becomes difficult or even impossible to assign accurate degrees of freedom for each testing statistic. In addition, the typical correction methods for multiple testing through spatial extent tend to be over-penalizing, and dichotomous decisions through thresholding under null hypothesis significance testing (NHST) are controversial in general and equally problematic in neuroimaging as well. For instance, the popular practice of only reporting “statistically significant” results in neuroimaging not only wastes data information, but also distorts the full results as well as perpetuates the reproducibility crisis because of the fact that the difference between a “significant” result and a “non-significant” one is not necessarily significant.

Here we propose a Bayesian multilevel (BML) framework for ISC data analysis that integrates all the spatial elements (i.e., regions of interest) into one model. By loosely constraining the regions through a weakly informative prior, BML conservatively pools the effect of each region toward the center, and improves collective fitting and overall model performance. The BML paradigm leverages the commonality or similarity among brain regions and the information across multiple levels embedded in the hierarchical data structure instead of leveraging the spatial extent adopted in the conventional correction method for multiple testing. In addition to potentially achieving a higher inference efficiency than the conventional LME approach, BML improves spatial specificity and easily allows the investigator to adopt a philosophy of full results reporting (instead of dichotomizing into “significant” and “non-significant” results), thus minimizing loss of information while enhancing reproducibility. A dataset of naturalistic scanning is utilized to illustrate the modeling approach with 268 parcels and to showcase the modeling capability, flexibility and advantages in reports reporting. The associated program will be available as part of the AFNI suite for general use.

## Introduction

Naturalistic scanning as an fMRI paradigm provides a window into shared brain responses at the population level under scenarios such as watching movies or listening to speech (Hasson et al., 2004, 2008). With minimal manipulation and dynamically evolving context, the naturalistic paradigm is closer to real life experiences, compared to typical task-related experiments while it is more engaging and less vulnerable to confounding effects such as head motions and physiological effects than resting-state scanning. It has been argued that, under a context closer to natural environment, neural responses are more reproducible and reliable than traditional simple repetitive stimuli (Hasson et al., 2010) due to the involvement of extensive cognitive processing (such as working memory, judgment, reasoning, social cognition, etc.). Its adoption has been steadily growing in investigating various aspects of brain functions such as music imagery (Zhang et al., 2017), early childhood development (Moraczewski et al., 2018), personality traits (Finn et al., 2018) and cognitive differences between schizophrenics and controls. For typical task-related designs, the focus is usually on identifying regions activated by an explicit task or condition. In contrast, the interest for naturalistic scanning data usually hinges on the synchronization or similarity between any pair of subjects. Specifically, one calculates the inter-subject correlation (ISC), which is the Pearson correlation between the EPI time series at the same voxel or region of the two subjects who underwent the same naturalistic-task scanning. In the end, an ISC value is obtained at each voxel or region for each subject pair, but the main issue is to summarize the results at the population level because of the complex relatedness among the subject pairs.

The current modeling approaches at the population level still face some challenges. Over the years, various methods including both parametric and nonparametric approaches have been developed to handle the complex relatedness (Bartels and Zeki, 2004; Hasson et al., 2008a; Wilson et al., 2008; Abrams et al., 2013; Kauppi et al., 2014; Schmälzle et al., 2013; Cantolon and Li, 2013; Schmälzle et al., 2015). For example, a popular but problematic approach is to first calculate the ISC value between a voxel’s BOLD time course of a subject and the average of that voxel’s BOLD time course among all other subjects (Kauppi et al., 2010; Honey et al., 2012; Schmälzle et al., 2013; Schmälzle et al., 2015), and then perform the typical group analysis (e.g., Student’s *t*-test) under the false assumption that all the ISC values are independent across subjects. Recently, we examined the validity of those methods, and proposed more rigorous approaches (Chen et al., 2016; Chen et al., 2017), among which the most flexible one in terms of analytical capability is linear mixed-effects (LME) modeling with a crossed random-effects structure (Chen et al., 2017). Nevertheless, there are a few limitations with the LME approach. (1) Input data redundancy. The ISC of each subject pair is used twice as input so that a balanced random-effects structure can be maintained in the LME model. (2) Difficulty of handling multiple testing. There are no similar counterparts of family-wise error (FWE) controllability available that are typically adopted in task-related experiments (e.g., cluster- or permutation-based methods). (3) Thorny issue of degrees of freedom. When an LME model is complicatedly structured, it becomes difficult or even impossible to assign accurate degrees of freedom for each testing statistic under the conventional null hypothesis significance testing (NHST). (4) Spatial specificity issue. The typical correction methods for multiple testing through spatial extent tend to dichotomize the statistical evidence and result in spatial clusters that are not necessarily aligned with anatomical structures in the brain, leading to interpretation ambiguities. (5) Model inefficiency. The methods of correction for multiplicity tend to be over-penalizing (Chen et al., 2019a), and dichotomous decisions under NHST through thresholding are controversial in general (McShane et al., 2017; Amrhein and Greenland, 2017) and equally problematic in neuroimaging as well (Chen et al., 2019a). For instance, the popular practice of only reporting “statistically significant” results in neuroimaging not only wastes data information, but also distorts the full results as well as perpetuates the reproducibility crisis because of the fact that the difference between a “significant” result and a “non-significant” one is not necessarily significant (Cox et al., 1977).

To address those limitations, here we propose a Bayesian multilevel (BML) framework that integrates all the spatial elements (i.e., regions of interest) into one model. Such a framework has been applied to typical task-related FMRI experiments (Chen et al., 2019a; Xiao et al., 2019) as well as matrix-based data analysis (Chen et al., 2019b; Yin et al., 2019). A dataset of naturalist scanning is utilized to illustrate the modeling approach and to showcase the modeling capability, flexibility and advantages in reporting results. This paper is a sequel (i.e., Part III) to our previous work of Part I (Chen et al., 2016) and Part II (Chen et al., 2017).

### Preamble

We summarize briefly the background, notations, framework, and structure of the ISC group analysis that were introduced in our previous work (Chen et al., 2016, Chen et al., 2017), since some shared concepts apply to the model formulation introduced here. Throughout this article, italic letters in lower case (e.g., *α*) stand for scalars; lowercase, boldfaced italic letters (***a***) and upper (***X***) cases for column vectors and matrices, respectively. With one group of *n* > 2 subjects *S*_1_, *S*_2_, …., *S_n_* and *m* spatial units (voxels or regions), the total number of unique ISC values per spatial unit is 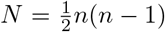. For the *k*th spatial unit (*k* = 1, 2, …, *m*), the ISC values {*r_ijk_*} correspond to *n* subject pairs (SPs), and they form a symmetric (*r_ijk_* = *r_jik_*, *i, j* = 1, 2, …, *n*) *n* × *n* positive semi-definite matrix 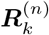 with diagonals *r_iik_* = 1 (Fig. 1, left). Their Fisher transformed version 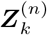 (Fig. 1, right) through *z* = arctanh(*r*) is usually adopted during analysis so that methods assuming Gaussian distribution may be utilized, as Fisher *z*-values are more likely to be Gaussian-distributed than raw Pearson correlation coefficients. The research of interest herein is focused on the population average effect of each region pair (*i*, *j*). Because 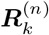 and 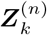 are both symmetric in (*i*, *j*), inferences at the population level can be made through the *N* elements in the lower triangular part (*i* > *j*, shaded gray in Fig. 1).

**Figure 1:**
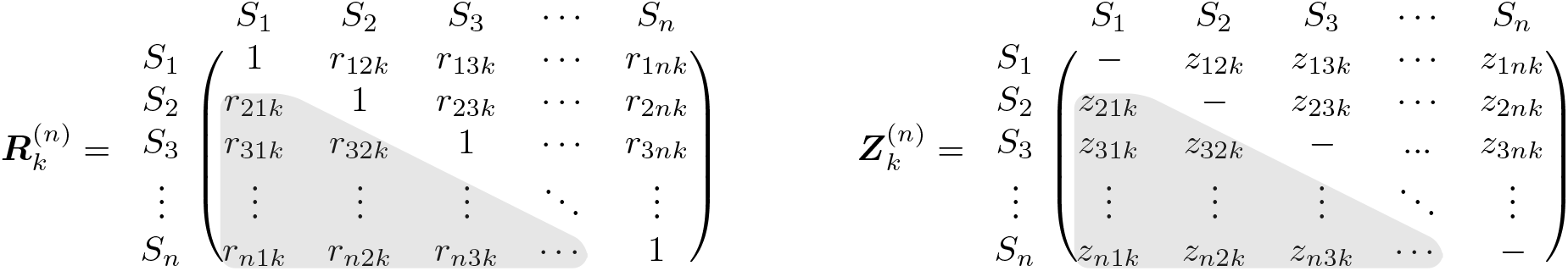
Inter-subject correlation (ISC) matrix 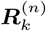 among the *n* subjects for the *k*th spatial unit and its Fisher-transformed counterpart 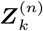. Due to the symmetry, only half of the off-diagonal elements (shaded in gray) are usually considered during ISC analysis.

The general interest of ISC analysis at the population level is the statistical inference about the population effect for each spatial unit. However, a complex issue to manage is that each ISC matrix element is correlated with some of others (Chen et al., 2017). Suppose that *z*_*i*_1_*j*_1_*k*_ and *z*_*i*_2_*j*_2_*k*_ are two *z*-values that are associated with the ISC values of the *k*th spatial unit, *r*_*i*_1_*j*_1_*k*_ and *r*_*i*_2_*j*_2_*k*_, of two SPs. When any pair of two elements in the ISC matrix, *z*_*i*_1_*j*_1_*k*_ and *z*_*i*_2_*j*_2_*k*_, involve four separate subjects (i.e., *i*_1_ ≠ *i*_2_ and *j*_1_ ≠ *j*_2_), we assume that the two elements are unrelated; that is, their correlation is 0. We denote the correlation between any two elements, *z*_*i*_1_*j*_1_*k*_ and *z*_*i*_2_*j*_2_*k*_, that pivot around a common subject (e.g., *i*_1_ = *i*_2_ or *j*_1_ = *j*_2_) as *ρ*, with the assumption that the relatedness *ρ* remains the same across all subjects. In other words, *ρ* characterizes the interrelatedness of *z*_*i*_1_*j*_1_*k*_ and *z*_*i*_2_*j*_2_*k*_ among the three subjects among which the two SPs share a common subject. To consider the group-wide set of ISCs, we further define ***z***_*k*_ = *vec*({*z_ijk_, i* > *j*}) to be the vector of length *N* whose elements are the column-stacking of the lower triangular part of the matrix ***Z***^(*n*)^ in Fig. 1. That is, ***z*** is the half-vectorization of 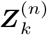 excluding the main (or principal) diagonal: 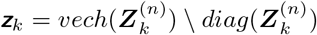. The variance-covariance matrix of ***z***_*k*_ can be expressed as the *N* × *N* matrix,

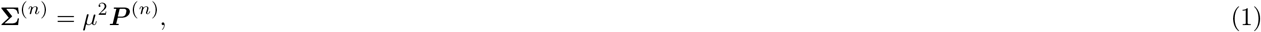

where *μ*^2^ is the variance of *z_ijk_, i* > *j*, and ***P***^(*n*)^ is the correlation matrix that is composed of 1 (diagonals), *ρ* and 0. An example of ***P***^(5)^ shown in Fig. 2. It has been analytically shown (Chen et al., 2016) that −1/[2(*m* – 2)] ≤ *ρ* ≤ 0.5 (when *m* > 3), and because of the presence of correlations among some elements of 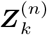, it becomes crucial to capture this correlation structure ***P***^(*n*)^ in any modeling framework.

**Figure 2:**
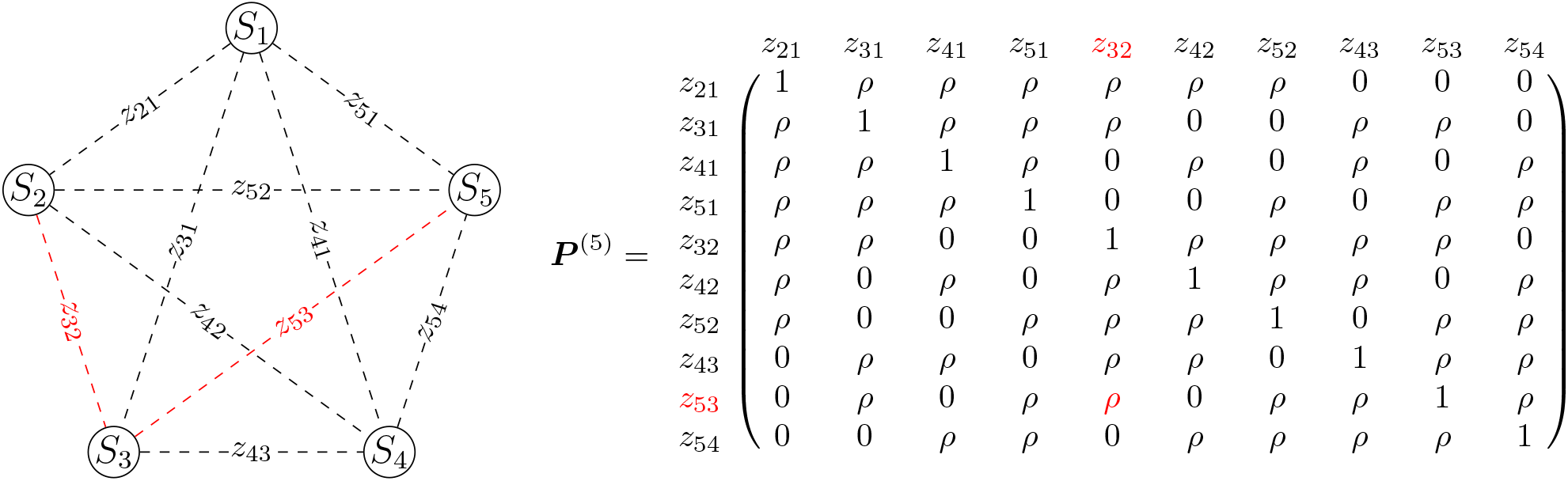
ISC with *m* = 5 subjects. Left: pictorial representation of 5 × 5 subject pairs (SPs). Right: The complex relatedness among the off-diagonal elements in 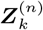 is illustrated with the correlation matrix ***P***^(5)^ for *n* = 5 subjects, in which *ρ* represents the correlation when two elements (e.g., *z*_32_ and *z*_53_, colored in red) are associated with a common subject (e.g., *S*_3_). Without loss of generality, the third index *k* in *z_ijk_* for brain location is dropped here for clarity.

The situation with two groups can be similarly formulated (Chen et al., 2016; Chen et al., 2017). Previously both nonparametric and parametric methods have been proposed to handle ISC analysis at the population level. Here we briefly summarize those methods, and lay out the background and motivations for our current work.

### ISC analysis with conventional approaches

In the early days, the pioneering work with naturalistic stimuli were conducted either within each subject when the natural stimulus was repeated several times (Hasson et al., 2008b) or through ISC for each SP separately without summarization at the group level (Hasson et al., 2004), in which case the ISC results were typically verified through seed-based correlation analysis (Hasson et al., 2004; Hasson et al., 2008b; Schmälzle et al., 2013). Later on, some investigators simply ran one-sample (Bartels and Zeki, 2004; Hasson et al., 2008a; Wilson et al., 2008; Abrams et al., 2013; Kauppi et al., 2014), two-sample (Schmälzle et al., 2013; Cantolon and Li, 2013) or paired (Abrams et al., 2013; Schmälzle et al., 2015) *t*-tests on Fisher-tranformed *z*-values {*z_ijk_*, *i* > *j*} of correlation coefficients, while it was generally acknowledged that the *N* elements {*z_ijk_*, *i* > j} were not independent, as illustrated in the correlation structure of ***P***^(*n*)^ in (1), thereby violating the independence assumption in the Student’s *t*-test and leading to the inflated degrees of freedom for the *t*-distribution as well as the underestimated standard error for the ISC estimate. The approach was mainly justified based on the observation that the null results generated by shifting each pair of time series by random steps roughly fitted to a *t*(*N* – 1)-distribution curve (Wilson et al., 2008).

Nonparametric methods have also been proposed in the previous ISC literature. For example, one popular approach with one group of subjects was to construct a null distribution for the whole brain by randomizing the time series across voxels and time points (e.g., circularly shifting each subject’s time series by a random lag so that they were no longer aligned in time across the subjects), as implemented into an analytical package ISC toolbox in Matlab (Kauppi et al., 2014; https://www.nitrc.org/projects/isc-toolbox/). One variation of this ISC approach is called leave-one-out: first calculate the ISC value of a subject between a voxel’s BOLD time course in the subject and the average of that voxel’s BOLD time course in the remaining subjects (Kauppi et al., 2010; Honey et al., 2012; Schmälzle et al., 2013; Schmälzle et al., 2015); then, perform Student’s *t*-test at the group level. However, a recent study has shown that all these methods led to largely inflated false positive rate (FPR) (Chen et al., 2016).

A new set of nonparametric approaches, based on subject-wise resampling at the population level, has been proposed recently (Chen et al., 2016). In addition to satisfying exchangeability and independence assumptions and accounting for the correlation structure in ***P***^(*n*)^, it was shown that proper FPR controllability under the conventional null hypothesis significance testing (NHST) can be achieved with subject-wise bootstrapping for ISC analysis with one group and with subject-wise permutation testing for the ISC comparison between two groups.

However, nonparametric methods are limited in terms of modeling flexibility. For instance, they have difficulty in incorporating explanatory variables; in addition, they are deficient, unwieldy and unconducive to data structure characterization and model comparisons. To counter these limitations, a linear mixed-effects (LME) modeling approach has been adopted (Chen et al., 2017) with the benefit that the LME platform offers wider adaptability, more powerful interpretations, and quality control checking capability than nonparametric methods. Specifically, the LME model with crossed random effects is applied with a data-doubling step that further conveniently tracks the subject index in easy implementations.

### ISC analysis with univariate linear mixed-effects modeling

Our previous work (Chen et al., 2017) adopts a linear-effects or multilevel model by decomposing an ISC effect *z_ijk_* into components associated subjects *i* and *j* at the *k*th voxel or region (*k* = 1, 2,…, *m*),

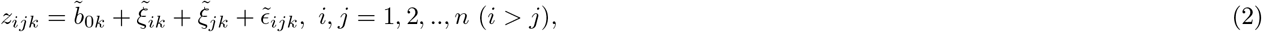

where 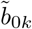 is the fixed effect (an unknown constant) under LME, representing the population ISC effect at the *k*th voxel or region; 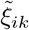 and 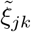 are additive and independent random effects attributable to subjects *i* and *j*, respectively, that are deviations from (or adjustments to) the population ISC effect 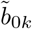; and 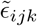 is the residual or error term for each SP (*i*, *j*). Due to the symmetric nature of the data structure in 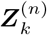, only half of the matrix elements excluding the diagonals (either the lower or upper triangular part of the matrix) are utilized in the model (2), and thus the index inequality of *i* > *j* is placed for the input data. As a special LME model, the formulation (2) can actually be conceptualized as a two-way random-effects ANOVA with the two subject-specific terms serving as random-effects factors. The two random effects 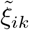 and 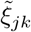 form a stratified structure, therefor the model (2) is sometimes referred to as a crossed (or cross-classified) structure with a factorial (or combinatorial) layout among the levels (or indices) *i* and *j* of the two subject-specific factors.

One important aspect of the LME framework in which nonparametric methods lack is that the interrelationships among the ISC values, as characterized in the correlation matrix ***P***^(*n*)^, can be quantitatively captured. With the assumption of independent Gaussian distributions, 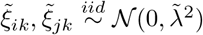 and 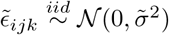, the model (2) can be solved under a two-way random-effects ANOVA or LME. A big advantage of the LME model (2) over the nonparametric methods is the capability of characterizing as well as maintaining the integrity of the data structure. For example, the extent of correlation *ρ*, as captured in ***P***^(*n*)^ of (1), between any two ISC effects that pivot around a common subject is related to the concept of intraclass correlation (ICC), and can be expressed as,

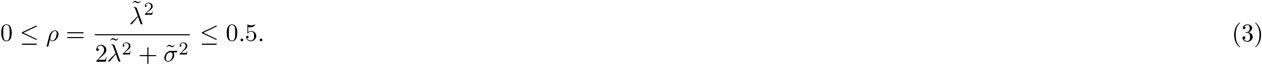

The LME model (2) can be easily extended to scenarios where the investigator would like to incorporate one or more subject-specific explanatory variables, either categorical (e.g., sex) or quantitative (e.g., age). For example, a model with one explanatory variable *x* can be formulated as,

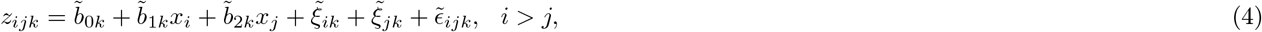

where the variable *x_i_* and *x_j_* are the values of the explanatory variable *x* for subjects *i* and *j*, respectively. Their corresponding effects 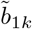 and 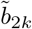 are presumably equal, but in the same vein as the practical implementation of subject-specific effects through two separate random-effects components as previously elaborated, the two fixed-effects components of 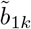 and 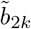 that are associated with the explanatory variable *x* would also have to be estimated separately through data duplication. The situation with more than one explanatory variable would be similar except for an expanded form, and this modeling strategy has been applied at the whole-brain voxel level to a few studies in the literature (e.g., Moraczewski et al., 2018; Finn et al., 2018).

Nevertheless, the LME framework faces two challenges. One challenge is input data redundancy. Even though the random-effects components, 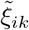 and 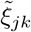, that are associated with the two subjects *i* and *j*, are assumed to follow the same Gaussian distribution 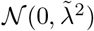 (that is why they are noted by the same symbol 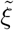), they would have to be treated as two separate random-effect components in practice when one solves the system through numerical implementations (e.g, function *lmer* in the *R* package *lme4*). Furthermore, due to the fact that only half of the off-diagonal elements in the matrix 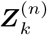 are utilized as input, the two random-effects components 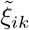 and 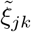 are generally not evenly arranged among all the SPs, leading to unequal estimation of the two random-effects components. On one hand, the two random-effects components 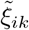 and 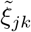 are basically cycled through those random effects from the *n* subjects, 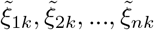, and the order of the two components, 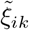 and 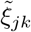, can be rearranged without any impact on the model formulation. On the other hand, balance cannot be achieved under all scenarios. For example, when *n* is odd, a balanced distribution between the two random-effects factors can be achieved through the following rearrangement: if the difference between the indices *i* and *j* is odd, switch their order (i.e., *z_ij_* effectively changes to *z_ji_*); otherwise, no change is made. However, when *n* is even, balance cannot be reached but can be approximated in the sense that the first index is alternatively one more (or less) than the second one^1^. Nevertheless, even if a balanced arrangement can be established between the two sets of indices (i.e., *n* is odd), simulations indicate unsatisfying FPR controllability for the population effect. Because of this limitation, a data doubling strategy (i.e., *i* ≠ *j*) was used with both the lower (*i* > *j*) and upper (*i* < *j*) triangular parts of the ISC matrix 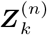 as input to achieve a balanced distribution between the two sets and proper FPR control (Chen et al., 2017). That is, with two random-effects components in the LME models, (2) and (4), two copies of the variance 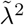 are estimated in the output. Also because of this data duplication, inferences have to be properly adjusted, compensating the inflated standard error (Chen et al., 2017).

The second challenge under the LME framework is multiplicity. It is worth noting that the LME model, (2) or (4), is analyzed through a massively univariate approach in which the same model is applied as many times as the number of voxels or regions. As in typical FMRI data analysis, the ISC analysis through LME at the whole brain or region level faces the issue of multiple testing. As the same LME model is applied to each voxel or region of interest (ROI) separately with the presumption that all the voxels or regions are isolated and unrelated. Such a modeling strategy would have to be followed by an extra step: paying the price of multiplicity for the false assumption, and one possible approach for the whole brain analysis is to control the overall FPR at the cluster level by leveraging the spatial extent among the neighboring voxels. First of all, permutation-based correction approaches would be impractical due to the prohibitively high computation cost. Even for cluster-based correction methods that are purely based on leveraging spatial extent (Monte Carlo simulations, random field theory), it remains challenging to estimate the spatial correlation due to the difficulty in separating the pure noise from the signal. Lastly, specific correction methods aside, the penalty is usually so severe that smaller brain regions may fail to survive the correction, in addition to other disadvantages of the massively univariate approach (Chen et al., 2019). If the LME approach is applied to a list of regions, Bonferroni correction would have to be employed because no spatial leveraging is obviously available, leading to even more severe penalty.

### Structure of the work

In light of the aforementioned backdrop, we believe that the univariate LME modeling approach at the whole brain level is inefficient because the common information shared across brain regions are fully ignored. Instead, we propose a more integrative and efficient approach, termed as Bayesian multilevel (BML) modeling, that could be used to confirm, complement or replace the LME method. As a first step, we adopt a group analysis strategy with LME by incorporating ROIs as a crossed random-effects component relative to each SP. Then we translate the LME model to the Bayesian framework through multilevel modeling on an ensemble of ROIs, and use this to resolve two issues that have been briefly mentioned: input data doubling and multiple testing. Those ROIs can be either determined independently from the current data at hand, or selected through various methods such as previous studies, an anatomical/functional atlas or parcellation. The proposed BML approach dissolves multiple testing through a multilevel model that more accurately accounts for data structure as well as shared information, and it consequentially improves inference efficiency.

The paper is structured as follows. In the next section, we first extend the region-wise LME model (2) to another LME by pivoting the ROIs as the levels of a random-effects factor, and then convert the extended LME model to a full BML. The BML framework does not make statistical inferences for each measuring entity (ROI in our context) in isolation. Instead, the BML weights and borrows the information based on the precision information across the full set of entities, striking a balance between data and prior knowledge; in a nutshell, the crucial feature here is that the ROIs, instead of being treated as isolated and unrelated with the univariate approaches, are associated with each other through a Gaussian-distribution assumption under BML. As a practical exemplar, we apply the modeling approach to an experimental ISC dataset with 68 subjects at 268 ROIs. In the Discussion section, we elaborate the advantages and limitations of BML modeling for ISC data analysis.

## Theory: ISC analysis through Bayesian multilevel modeling

Herein Roman and Greek letters are used, respectively, to differentiate fixed and random effects in the conventional statistics context such as ANOVA and LME on the righthand side of a model equation. Although the terms of “fixed” and “random” effects are genuinely non-Bayesian, we still use them here as we expect most readers to be familiar with the conventional terminology. For instance, a conventional fixed-effects parameter under ANOVA and LME is treated as constant that is shared by all entities (e.g, subjects, ROIs), and a random-effect parameter as variable because it differs from one entity (e.g., subject, ROI) to another. The conventional distinction of fixed-vs. random-effects is replaced by one that separates the modeling decision (a parameter as varying or non-varying) under the Bayesian framework from the inference decision (e.g., prior choices or partial pooling) (Gelman, 2005).

### Bayesian modeling based on three-way random-effects ANOVA

We start with with the simple LME model (2), without the complication of explanatory variables, for ISC analysis at *m* ROIs in the brain instead of whole brain voxel-wise modeling. With the Gaussian-distribution assumptions for 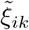, 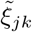, and 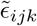, the *m* univariate LME models in (2) can be solved independently, but for the sake of model comparisons, the *m* separate LMEs can be merged into one LME by pooling the residual variances across the *m* ROIs with the ROI index *k* incorporated into the conventional LME formulation (2),

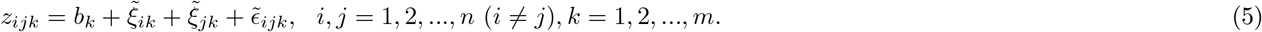

The two approaches, (2) and (5), usually render similar inferences unless the sampling variances are dramatically different across the *m* ROIs. To compare different models through information criteria (Vehtari et al., 2017), we can solve the LME (5) in a Bayesian fashion,

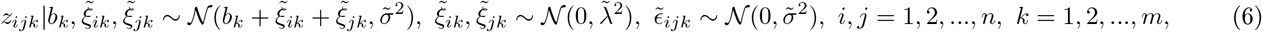

where the effects *b_k_* are assigned with a noninformative prior (i.e., uniform distribution) so that no information is shared among the ROIs, leading to virtually identical inferences as the LME (5). In fact, all the three LME models, (2), (5), or (6), share the same feature: they do not involve any information pooling among the ROIs in the sense that the information at one ROI is assumed to reveal nothing about any other ROIs. Therefore, these three LME models all face the same issue of multiplicity and may potentially lead to overfitting.

To improve model fitting and achieve higher efficiency, we first adopt a three-way random-effects ANOVA or LME model by adding ROIs as random effects, and formulate the following platform with data from *n* subjects,

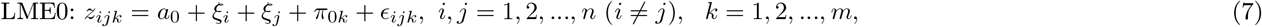

where *a*_0_ represents the population ISC effect across all ROIs and all subjects; *ξ_i_* and *ξ_j_* code the deviation or random effect of the *i*th and *j*th subject from the overall mean *b*_0_, respectively, and both share the same presumed *iid* Gaussian distribution 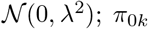 embodies the random effect (or the deviation relative to the population effect *a*_0_) at the *k*th ROI, and is assumed to be *iid* with 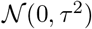; and *ϵ_ij_* is the residual term that is assumed to follow 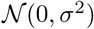. Put in a different way, under the LME model (7), each data point *z_ijk_* is disentangled as the superimposition of three random-effects components, two (*ξ_i_* and *ξ_j_*) for each SP and one (*π*_0*k*_) for each region. The tilde notation above the parameters in the previous four LME models, (2), (4), (5) and (6), is removed hereafter due to the inclusion of ROIs as random effects in the expanded models such as (7) as well as the removal of subscript for the ROI index *k* among some parameters (e.g., *ξ_i_* and *ξ_j_* in the model (7)).

Under the extended LME model (7), the correlation between two SPs, (*i*_1_, *j*) and (*i*_2_, *j*) (*i*_1_ ≠ *i*_2_), that share a common subject *S_j_* can be derived as,

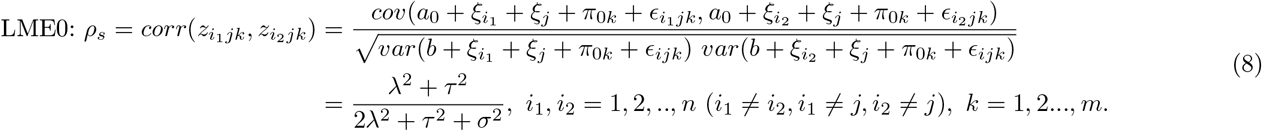

Similarly, the correlation between two ROIs, *k*_1_ and *k*_2_, due to the fact that they are measured from the same SPs, can be derived as,

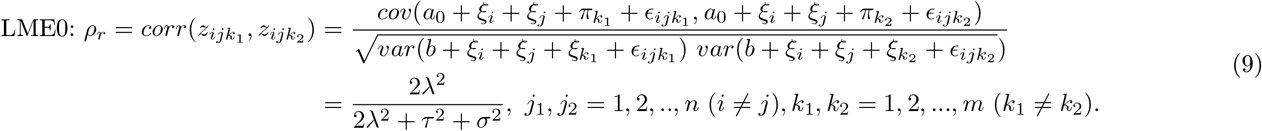

Due to the incorporation of ROI effects into the extended LME model (7), a slightly different formulation (8) at the group level for the correlation between two SPs that share a common subject exists from the interrelationship (3) at the individual subject level. Because of this difference, the upper bound of 0.5 in (3) does not hold for *ρ_s_* in (8) and is replaced by 1.

In addition to the challenge of input data redundancy discussed in the Introduction, now we have a different hurdle in place of multiplicity. Under this new LME framework (7), we need to refocus on the effects of interest. The overall ISC effect *a*_0_ across all ROIs is usually not our focus; instead, it is the ISC effect each ROI,

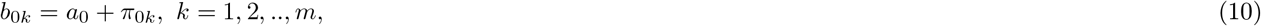

that is typically of research interest. However, the LME formulation or three-way random-effects ANOVA (7) cannot offer a solution in making inferences regarding the ROI effects *b*_0*k*_: to estimate *b*_0*k*_, the LME (7) would become over-parameterized or overfitting.

To proceed, a paradigm shift is needed here. We adopt a Bayesian approach similar to, but extended the LME model (2) from, our previous work for ROI-based group analysis for neuroimaging data (Chen et al., 2019a) as well as the BML approach for matrix-based analysis (Chen et al., 2019b),

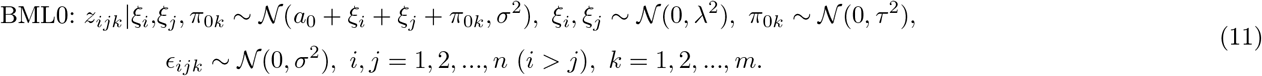

In fact, the effect decomposition of *z_ijk_* under the BML framework (11) is basically the same as its LME counterpart (7). The different model expression here is formulated to accentuate the paradigm shift and to emphasize the fact that the responses *z_ijk_* under BML are conditional on the parameters and priors.

Both of the aforementioned challenges under the LME model (2) can be resolved now under the BML framework (11). First, only half of the off-diagonal elements (e.g., the lower triangular part) in ***Z***^(*n*)^ are required as input under BML through a numerical implementation of multi-membership modeling scheme (Bürkner, 2018). Second, with a prior (e.g., noninformative uniform distribution) for *a*_0_, the posterior distribution for each ROI can be obtained through the formulation (10). In addition, the ISC effect that is attributable to each subject can be similarly derived through the corresponding posterior distribution with

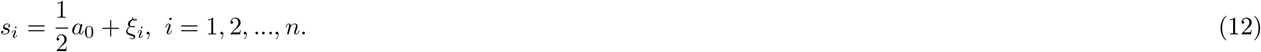

The factor of 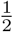 in the subject-specific effect formula for *s_i_* in (12) reflects the fact that the effect of each SP is evenly shared between the two associated subjects. The subject-specific effects *s_i_* can be utilized to assess the contribution or importance of a subject relative to all other subjects, which might provide some auxiliary information for further association with, for example, subject-level effects such as sex, disease, age or behavioral data.

Recently we applied the BML modeling approach to matrix-based analyses (Chen et al., 2019b) when the input data are either functional (e.g. inter-region correlation) or structural (e.g., white matter properties among gray matter regions) attribute matrix from each subject. In that case, the intricacy lies in the interrelationships among the brain region pairs while the summarization or generalization hinges upon the subjects, and three basic entity-level components are specified in the corresponding BML model: subject and the two regions that are associated with each region pair. In contrast, ISC analyses deal with the interrelationships among SPs while at the same time the summarization or generalization is made across subjects; the regions under BML are pooled together among each other through the shrinkage effect of the Gaussian distribution (Chen et al., 2019a). In fact, the theoretical aspects of BML application for ISC analyses can largely be borrowed from our previous work for MBA (Chen et al., 2018b) by swapping the entities between subject and region.

### Further extensions of BML for ISC analyses

The LME0 model in (7) can be expanded or generalized by including two types of random-effects interaction components: one component is the SP-specific term (i.e., the interaction between two subjects), and the other component is the interaction between a region and a subject. The expansions lead to three more LME models, corresponding to three different combinations of the two extra effects, as shown below,

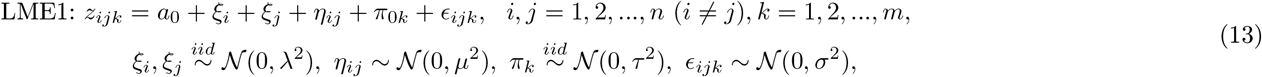

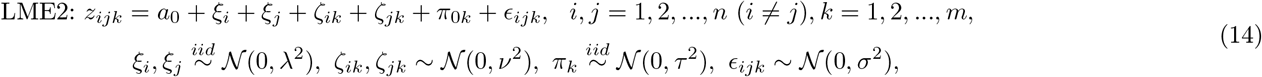

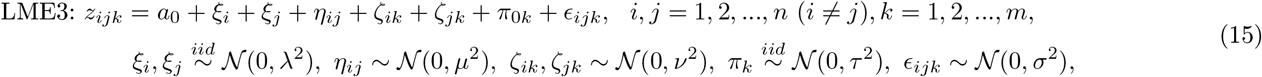

where *η_j_* is the effect of the SP that is associated with subjects *i* and *j* (i.e., the interaction effect between two subjects *i* and *j*) relative to the overall effect *a*_0_ and the two subject effects, *ξ_i_* and *ξ_j_*, while *ζ_ik_* and *ζ_jk_* are the interaction effects between subject *i* and region *k* as well as the interaction between subject *j* and region *k*, respectively. We note that the SP-specific effect *η_ij_* captures the unique effect of each SP in addition to the overall effect *a*_0_ and the common effects from the two involved subjects, *ξ_i_* and *ξ_j_*; the same subtlety applies to the subject-region interactions *ζ_ik_* and *ζ_jk_*.

The two ICC measures in (8) and (9) can be correspondingly updated to the following for the three LME models,

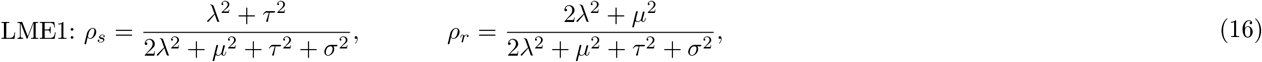

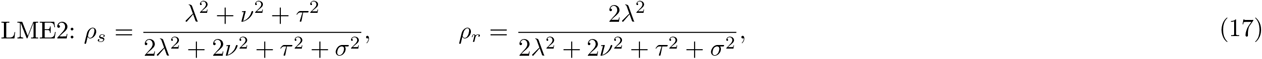

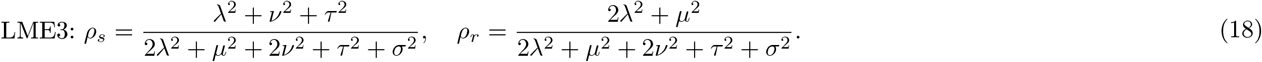

Among the four LME models, LME0 is the simplest and LME3 is the most complex and inclusive, while LME1 and LME2 are intermediate. Traditionally, these LME models can be compared based on the tradeoff between model performance and complexity (e.g., number of parameters), with likelihood ratio testing and information criteria such as Akaike information criterion (AIC) and the so-called Bayesian information criterion (BIC) (Bates et al., 2015). As the number of components in a model increases, so does the number of parameters to be estimated. For example, with *n*(*n* – 1)*m* data points *z_ijk_* as input, the total number of parameters involved at the right-hand side of the model LME3 in (15) is *n*(*n* – 1) + 2*mn* + 2*n* + 1. For the model LME3 to be identifiable, it is a prerequisite that the following relationship be met,

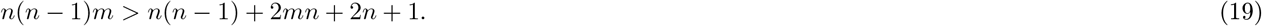

When there are two or more ROIs (*m* > 1), the prerequisite (19) means that, to prevent LME3 from being overparameterized, a condition for the number of subjects, derived from a quadratic form of *n*, is 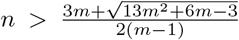. Such a lower bound for *n* is a decreasing function of *m* as shown in a few instances: when *m* = 2, 5,10,100, and 1000, *n* > 7,4, 3, 3, and 3, respectively. For the special case of one region (*m* = 1), all the three models, LME1, LME2, and LME3, reduce to LME0 in which the SP-specific effects *η_j_* and the interaction effects between subjects and regions, *ζ_ik_* and *ζ_jk_*, cannot be differentiated from the residuals *ϵ_ijk_* and region-specific effects *π*_0*k*_, respectively.

We further consider two types of BML extension based on the primary model BML0 in (11). The first type involves potential interaction effects, in parallel with the three LME expansions from LME0. Specifically, by incorporating the interaction effect between the two regions of each RP as well as the interaction effect between each region and each subject, we have three more BML models (corresponding to the LME models of the same index):

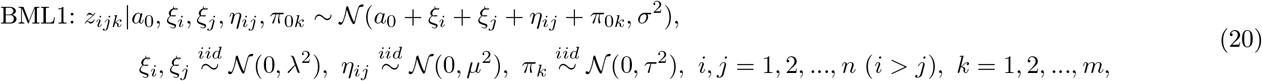

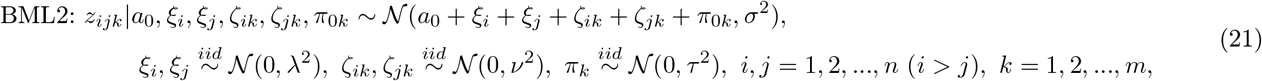

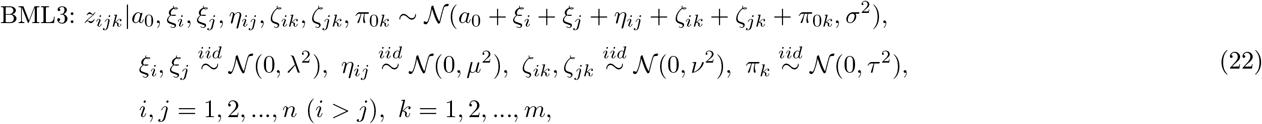

where *η_j_* is the SP-specific effect or the interaction between subjects *i* and *j*, while *ζ_ik_* is the interaction effect between subject *i* and region *k* and *ζ_jk_*, between subject *j* and region *k*. The two interaction effects, *ζ_ik_* and *ζ_jk_*, are considered as two members, *i* and *j*, of a multi-membership cluster. Because of the sheer number of parameters, their LME counterparts are not always identifiable (e.g., because the prerequisite (19) is violated), but these BML models can be analyzed under the Bayesian scheme because of the constraints and regularization applied through priors. Similar to the LME case, complexity increases from BML0 to BML3. Under these three extended BML models, the region- and subject-specific effects can be similarly reassembled through (10) and (12), respectively.

### Inclusion of explanatory variables under BML for ISC analyses

Another type of model extension is to investigate the effect associated with a subject-level (e.g., sex, disease, genotype, age, behavioral measure) explanatory variable. For example, with one explanatory variable, BML0 (11) and BML1 (20) can be directly expanded by adding a covariate *x*,

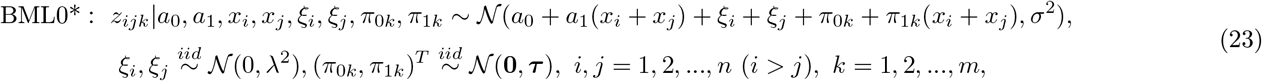

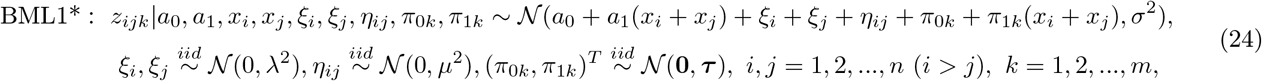

where *τ* is a 2 × 2 variance-covariance matrices.

Five aspects are noteworthy about the two extended models, BML0* and BML1*. First, multi-membership modeling allows us to utilize only half of the off-diagonals in the ISC matrix from each subject as input, as indicated by the index relationship *i* > *j*. Second, the effect associated with the covariate *x* at the population level, *a*_1_, and at the region level, *π*_1*k*_, is shared by all subjects (including SPs), thus a simplified notation for a derived covariate 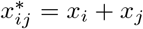 for each SP can be adopted for easier numerical implementation. This is in contrast to the LME counterpart in which two separate effects would have to be included in the model. Third, the inclusion of any subject-level explanatory variable in the model is intended to account for cross-subject variation in the data, thereby precluding the justification for incorporating the subject-region interaction effects, *ζ_ij_* and *ζ_jk_*, as shown in BML2 (21) and BML3 (22). It light of this consideration, we do not consider any extended models, in the presence of any subject-specific covariate, that correspond to BML2 (21) and BML3 (22). Four, cases with more than one explanatory variable can be similarly formulated as in the BML0* and BML1*. Lastly, under BML0* or BML1*, the region- and subject-specific effects can be similarly reassembled through (10) and (12), respectively; in addition, the region-specific effect for the covariate *x* can be derived through,

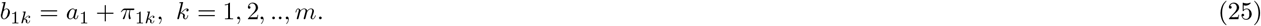

To recapitulate our modeling strategy here about ISC analyses, we first untangle each SP-specific effect into the additive effects of the two involved subjects through a multi-membership structure. The partition of ISC effect allows us to maintain the relatedness structure as embodied in the correlation matrix ***P***^(*n*)^. Because of this untangling step, we can obtain the relative contribution, *s_i_* in (12), from each subject even though the input data (ISC values) are the jointed contributions from SPs, not individual subjects. In addition, the cross-region effects (and sometimes subject-region interaction effects) are included in the BML model to account for cross-region variability. The main difference between univariate LME (Chen et al., 2017) and BML lies in the assumption about the brain regions: the effects (e.g., *π*_0*k*_ and *π*_1*k*_ in (24)) are assigned with a Gaussian prior under BML while they are assumed to have a noninformative flat prior under the corresponding LME model. In other words, the effect at each region is estimated independently from other regions under univariate LME, thus there is no information shared across regions. In contrast, the effects across regions are shared, regularized and partially pooled through the Gaussian assumption under BML for the effects across regions; the Gaussian assumption about cross-region variability shares the same rationale as the cross-subject Gaussian distribution. On one hand, it is this pooling effect that drags the region effects from both ends toward the center, resulting in conservative inferences relative to univariate LME before taking into consideration any correction for multiple testing. On the other hand, it is partial pooling that allows us to have an integrative model that sidesteps the multiplicity issue.

### Implementations of BML for ISC analyses

With the layout of three or more crossed random-effects components under the LME models such as LME0 (7), LME1 (13), LME2 (14), and LME3 (15), we would have to duplicate the input data and utilize both the lower (*i* > *j*) and upper (*i* < *j*) triangular parts of the ISC matrix 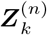 as input so that a balanced data structure (*i* ≠ *j*) can be maintained for the practical reason of typical numerical implementations, as adopted in our previous work for whole-brain voxel-wise ISC analyses (Chen et al., 2017) using the function *lmer* in R package *lme4* (Bates et al., 2015). Similarly, each explanatory variable (e.g., sex, behavioral measure) has to be incorporated inot the LME model with two copies, one for each subject within each SP. In doing so, the uncertainty estimates (e.g., standard deviations) for the obtained effect estimates need to be adjusted for artificially doubling the data. In addition, each of the effect pairs such as *ξ_i_* and *ξ_j_* in the four LME models above as well as *ζ_ik_* and *ζ_jk_* in LME2 and LME3, are treated as two separate random-effects components in real implementations even though they are the same effect within each component.

For the BML systems such as BML0 to BML3 and their counterparts with covariates, we borrow the terminology and implementation strategy from multi-membership modeling. Specifically, we consider both subject-specific effects of *ξ_i_* and *ξ_j_* as two members (samples or substantiations) from the same set of parameters (or the same list of subjects in the neuroimaging context) with equal weights of 1, reducing the number of associated parameters from 2*n* in the LME counterpart to *n* and maintaining the original index constraint *i* > *j* in the BML models here. Substantial runtime can be saved through the avoidance of input data redundancy.

As no analytical solution is available for BML models in general, we draw samples from the posterior distributions via Markov chain Monte Carlo (MCMC) simulations with the algorithms implemented in Stan, a publicly available probabilistic programming language and a math library in C++ on which the language depends (Stan Development Team, 2019). In Stan, the main engine for Bayesian inferences is adaptive Hamiltonian Monte Carlo (HMC) under the category of gradient-based MCMC algorithms (Betancourt, 2018). The present implementations are executed with the *R* package *brms* that is is based on Stan, and multi-membership modeling is directly available in *brms* (Bürkner, 2017; Bürkner, 2018).

For typical BML models, examples of each of the priors (e.g., hierarchical Gaussian distributions) for cross-region and cross-subject effects as well as their interactions have been laid out in the previous section. For example, we adopt an improper flat (noninformative uniform) distribution over the real domain or a weakly informative distribution such as Cauchy or Gaussian for population parameters (e.g., *a*_0_ and *a*_1_ in BML0* (23) and BML1* (24)), depending on the minimal requirement to cope with the amount of data present; in other words, one may adopt a noninformative prior if a large amount of information is available in the data at the population level. As for assigning hyperpriors, we follow the general recommendations in Stan (Stan Development Team, 2019). Specifically, for the scaling parameters at the region and subject level, the standard deviations for the cross-region and cross-subject effects, *ξ_i_*, *ξ_j_*, and *π_k_* as well as their interactions, we adopt a weakly informative prior such as a Student’s half-*t*(3,0,1)^2^ or half-Gaussian 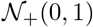 (restricting to the positive values of the respective distribution). For covariance structure (e.g., *τ* in BML0* (23) and BML1* (24)), the LKJ correlation prior^3^ is used with the shape parameter taking the value of 1 (i.e., jointly uniform over all correlation matrices of the respective dimension) (Gelman et al., 2017). Lastly, the standard deviation σ for the residuals is assigned using a half Cauchy prior with a scale parameter depending on the standard deviation of *z_ijk_*. To summarize, besides the Bayesian framework under which hyperpriors provide a computational convenience through numerical regularization, the major difference between BML and its univariate GLM counterpart is the application of the Gaussian prior in the BML models that play the pivotal role of pooling and sharing the information among the brain regions. It is this partial pooling that effectively takes advantage of the effect similarities among the ROIs and achieves higher modeling efficiency.

Bayesian inferences are usually expressed in terms of the whole posterior distribution of each effect of interest. For practical considerations in results reporting, point estimates from these distributions such as mean and median are typically used to show the effect centrality, while quantile-based (e.g., 90%, 95%) intervals or highest posterior density intervals also provide a condensed and practically useful summary of the posterior distribution. A typical workflow to obtain the posterior distribution for an effect of interest is the following. Multiple (e.g., 4) Markov chains are usually run in parallel with each of them going through a predetermined number (e.g., 2000) of iterations, half of which are thrown away as warm-up (or “burn-in”) iterations while the rest are used as random draws from which posterior distributions are derived. To gauge the consistency of an ensemble of Markov chains, the split 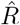 statistic (Gelman et al., 2014) is provided as a potential scale reduction factor on split chains and as a diagnostic parameter to assist the analyst in assessing the quality of the chains. Ideally, fully converged chains correspond to 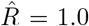, but in practice 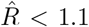 is considered acceptable. Another useful statistic, effective sample size (ESS), measures the number of independent draws from the posterior distribution that would be expected to produce the same amount of information of the posterior distribution as is calculated from the dependent draws obtained by the MCMC algorithm. As the sampling draws are not always independent of each other, especially when MCMC chains mix slowly, one should make sure that the ESS is large enough (e.g., 200) so that a reasonable accuracy can be achieved to derive the quantile intervals for the posterior distribution. With the number of cores equal to or large than the number of MCMC chains, the typical BML analysis can be effectively conducted on any system with at least 4 CPUs.

## BML applied to ISC data

To demonstrate the modeling capability and performances of BML, we used a dataset from the Child Mind Institute Healthy Brain Network (CMI-HBN), a publicly available naturalistic scanning dataset (Alexander et al. 2017). Briefly, the dataset consisted of a community-based sample of generally healthy children and adolescents who were scanned while resting as well as watching two different videos. Rich phenotypic data are also available for each individual. We focus here on data acquired during “The Present,” an animated short about a boy who receives a puppy as a gift. The video has a social theme and is emotionally evocative, which led us to hypothesize that it would evince individual differences along a phenotypic spectrum related to social functioning. Data used here come from the CMI-HBN data releases 1 and 2, which represented all of the available data in January 2018 when we began the project.

Functional MR images were acquired with the following EPI scan parameters: B0 = 3 T, flip angle = 31°, TR = 800 msec, TE = 30 msec, 60 slices, voxel size = 2.4 mm isotropic, multiband factor = 6, 250 volumes with a total scanning time of 3:20 min:sec. Other details, including parameters for anatomical scans as well as full protocols for MRI and phenotypic data, can be found in the data descriptor (Alexander et al. 2017) and at the following URL: http://fcon_1000.projects.nitrc.org/indi/cmi_healthy_brain_network/

Data were preprocessed as follows. First, we used Freesurfer (Fischl, 2012) to extract subject-specific ventricle and white-matter masks using each subject’s anatomical image. Next, we used the afni_proc.py program in AFNI (Cox, 1996) to perform the following preprocessing steps on the functional images: despiking, head motion correction, affine alignment with anatomy, nonlinear alignment to a standard template, and smoothing with an isotropic FWHM of 5 mm. Confounding effects during preprocessing included: the first three principal components of the ventricles, local white matter regressors generated from fast ANATICOR (Jo et al, 2010), each subject’s 6 motion time series, their derivatives and linear polynomials for slow drifts. Censoring of time points was performed whenever the per-time-point motion (Euclidean norm of the motion derivatives) was 0.3 mm or more or when more than 10% of the brain voxels were outliers. Censored time points were set to zero rather than removed altogether (this is the conventional way to do censoring, but especially important for inter-subject correlation analyses, to preserve the temporal structure across participants). Because this is a pediatric sample, we used a recently developed pediatric template brain as the standard template (“Haskins template”; Molfese et al., in prep).

Our primary phenotypic measure of interest was the Social Responsiveness Scale-2, abbreviated here as SRS (Constantino and Gruber, 2012). This parent-report scale measures the presence and severity of social impairment using items such as “seems much more fidgety in social situations than when alone”, “takes things too literally and doesn’t get the real meaning of a conversation”, and “avoids eye contact or has unusual eye contact”. There are 65 total items and each is rated on a Likert scale from 0-3; higher scores indicate poorer social functioning.

We selected a subset of subjects for analysis based on the following criteria: (1) a usable T1-weighed anatomical image (for registration purposes), (2) the functional movie-watching run of interest (“The Present”), with at least 85% (213/250) volumes remaining after censoring of head motion and outliers, (3) valid demographic information including age and sex; and (4) a valid SRS score. There were 68 subjects that met these criteria (age range = 6-17 years, mean ± standard deviation = 10.8 ± 3.1 years; 30 females). SRS scores followed a right-skewed distribution with range = 3-140, median (mean) = 43.5 (53.3), and median absolute deviation (standard deviation) = 17 (33.6). In this subset, there was negligible correlation between age and SRS (*r* = 0.046) or between head motion (as measured by mean frame-wise displacement) and SRS (*r* = −0.064). There was a moderate negative correlation between age and head motion (*r* = −0.25). Males and females did not differ much in age (males 10.35 ± 2.95 years, females 11.3 ± 3.19 years). However, SRS scores were moderately higher among males than females (males 58.26 ± 35.26, females 47.07 ± 30.88).

Owing to the computational intractability of conducting BML at the voxel-wise level, we defined ROIs using a preexisting functional brain parcellation (Shen et al., 2013), which contains 268 regions covering the whole brain (cortex, subcortex and cerebellum). It was originally defined in MNI space and nonlinearly warped to Haskins template space for purposes of this study. Region-wise time courses for each subject were calculated by averaging the signal of all the voxels in each region at each time point. Thus, the final dataset that entered into the ISC calculation consisted of 268 regions × 250 time-points × 68 subjects. To demonstrate that the method is robust to the choice of ROIs and spatial resolution of the parcellation, we also conducted the same analysis using a coarser, anatomically defined parcellation containing 107 nodes that is included as part of the Haskins template space (Molfese et al., in prep).

The ISC data of Fisher-transformed *z*-values from the *n* = 68 subjects at *m* = 268 ROIs were analyzed with three models: BML0* (23) and BML1* (24), and the region-wise LME model that corresponds to BML1*. Three explanatory variables (SRS, Age, and Sex), plus their two- and three-way interactions, leading to a total of eight effects of interest at each ROI: overall ISC (intercept), main effects (SRS, Age, Sex), two-way interactions (SRS:Age, SRS:Sex, Age:Sex), and three-way interaction (SRS:Age:Sex). The ROI dataset was analyzed with the three models using the R package *brms*. Runtime for BML was three weeks on a Linux system of Fedora 25 with AMD Opteron 6376 at 1.4 GHz; in contrast, the runtime of the same model with the coarser parcellation of 107 ROIs was five days.

To compare the two BML models, we assessed their point-wise out-of-sample prediction accuracy through the leave-one-out information criterion (LOOIC). As the LOOIC for the BML1* model (with subject pair specific effects) relative to BML0* (without subject pair specific effects) is −56406.34 ± 474.65, the higher predictive accuracy of BML1* is shown by its substantially lower LOOIC than BML0*. We thereafter focus our results discussion on BML1*.

The summary of the BML1* parameter estimates is shown in Table 1. One noteworthy aspect is that the interaction effect *η_ij_* of subject pairs was substantial with a standard deviation λ = 0.091 (with a 95% quantile interval of [0.090,0.092], Table 1), and such an interaction was stronger than the additive effects of individual subjects *ξ_i_* or *ξ_j_* with a standard deviation *μ* = 0.079 (with a 95% quantile interval of [0.076, 0.084], Table 1). In other words, cross-subject-pairs effects *η_ij_* account for a little more ISC variability than cross-subjects effects *ξ_i_* and *ξ_j_*. These results justify our adoption of the extended BML1* model (24) that contains the cross-subject-pairs effects *η_ij_* instead of BML0* (23) without the effect *η_ij_*. This result is also interesting from a scientific perspective, as it suggests that the interaction between a given subject pair is more important for determining ISC levels than either of the two subjects on their own. In other words, it is generally *not* the case that an individual subject tends to have high (or low) ISC values across the board (i.e., with all potential pairs); rather, it is the specific subject pair that explains more variability in observed ISC effects.

**Table 1:** Summary results from the ISC dataset fitted with an extended version of BML1* in (24) and its LME counterpart. The column headers Estimate, SD, QI, and ESS are short for effect estimate, standard deviation, quantile interval, effective sample size, respectively. LME1* shares the same effect components as BML1*, and shows virtually the same effect estimate for the population mean *b*_0_ and the standard deviations for those effect components despite: (1) the two modeling frameworks were solved through two different numerical schemes (REML for LME and MCMC for BML); and 2) in practice the input data for LME3 had to be duplicated to maintain the balance between the two crossed random-effects components associated with each subject pair. In addition, the nearly identical parameter estimates between the two models indicate that the use of priors under BML had a negligible effect. However, the LME framework cannot provide uncertainty measures for those variances, as indicated by the dashes in the table. 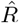 is the split statistic of a convergence indicator for the Markov chains. All 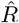 values under BML1* were less than 1.1, indicating that all the four MCMC chains converged well. The effective sample sizes (ESSs) for the population- and region-level effects were large enough to warrant quantile accuracy in summarizing the posterior distributions for region-specific effects. The correlations among the eight cross-region effects under BML are not shown in the table because their inferences are not available under LME.

| Term | BML1* |  |  |  |  | LME1* |  |
| --- | --- | --- | --- | --- | --- | --- | --- |
| | Estimate | SD | 95% QI | ESS | $\hat{R}$ | Estimate | SD |
| population-level effects |  |  |  |  |  |  |  |
| $a_0$: Intercept | 0.057 | 0.064 | [0.045, 0.069] | 104 | 1.04 | 0.057 | 0.063 |
| $a_1$: SRS | -1.27e-4 | 8.35e-5 | [-2.86e-4, 3.99e-5] | 492 | 1.00 | -1.31e-4 | 5.54e-5 |
| $a_2$: Age | -1.12e-3 | 8.78e-4 | [-2.78e-3, 5.80e-4] | 377 | 1.00 | -1.14e-3 | 6.06e-4 |
| $a_3$: Sex | -3.54e-3 | 5.38e-3 | [-1.39e-2, 7.37e-3] | 349 | 1.01 | -3.70e-3 | 3.76e-3 |
| $a_4$: Age:Sex | 7.85e-4 | 3.23e-4 | [ 1.51e-4, 1.42e-3] | 317 | 1.01 | 7.54e-4 | 2.44e-4 |
| $a_5$: SRS:Sex | 9.22e-6 | 3.07e-5 | [-5.12e-5, 7.04e-5] | 244 | 1.01 | 9.45e-6 | 2.21e-5 |
| $a_6$: Age:SRS | 5.53e-6 | 5.35e-6 | [-4.56e-6, 1.64e-5] | 295 | 1.00 | 5.68e-6 | 3.81e-6 |
| $a_7$: Sex:Age:SRS | 1.54e-6 | 6.30e-6 | [-1.06e-5, 1.42e-5] | 257 | 1.01 | 1.75e-6 | 4.56e-6 |
| cross-subjects effects (levels: 68) |  |  |  |  |  |  |  |
| $\lambda$: standard deviation for $\xi_i, \xi_j$ | 0.079 | 0.060 | [ 0.076, 0.084] | 561 | 1.01 | 0.079 | - |
| cross-subject-pairs effects (levels: 2278) |  |  |  |  |  |  |  |
| $\mu$: SD for $\eta_{ij}$ | 0.091 | 0.058 | [ 0.090, 0.092] | 395 | 1.01 | 0.091 | - |
| cross-ROIs effects (levels: 268) |  |  |  |  |  |  |  |
| $\tau_0$: SD for Intercept $\pi_{0k}$ | 0.106 | 0.060 | [ 0.102, 0.111] | 66 | 1.06 | 0.106 | - |
| $\tau_1$: SD for SRS $\pi_{1k}$ | 1.19e-4 | 6.14e-6 | [ 1.08e-4, 1.31e-4] | 563 | 1.01 | 1.23e-4 | - |
| $\tau_2$: SD for Age $\pi_{2k}$ | 1.40e-3 | 7.0e-5 | [ 1.27e-3, 1.54e-3] | 705 | 1.01 | 1.44e-3 | - |
| $\tau_3$: SD for Sex $\pi_{3k}$ | 8.0e-3 | 4.0e-4 | [ 7.22e-3, 8.80e-3] | 948 | 1.00 | 8.22e-3 | - |
| $\tau_4$: SD for Age:Sex $\pi_{4k}$ | 1.27e-3 | 7.54e-5 | [ 1.13e-3, 1.43e-3] | 1442 | 1.00 | 1.36e-3 | - |
| $\tau_5$: SD for SRS:Sex $\pi_{5k}$ | 1.05e-4 | 6.50e-6 | [ 9.29e-5, 1.18e-4] | 1468 | 1.00 | 1.14e-4 | - |
| $\tau_6$: SD for Age:SRS $\pi_{6k}$ | 2.03e-5 | 1.20e-6 | [ 1.79e-5, 2.27e-5] | 1130 | 1.00 | 2.22e-5 | - |
| $\tau_7$: SD for Sex:Age:SRS $\pi_{7k}$ | 1.48e-5 | 1.62e-6 | [ 1.16e-5, 1.79e-5] | 1274 | 1.00 | 1.96e-5 | - |
| residuals |  |  |  |  |  |  |  |
| $\sigma$: SD for residuals | 0.160 | 0.058 | [ 0.160, 0.160] | 3097 | 1.00 | 0.160 | - |

The eight effects of interest can be shown with their respective posterior distributions. However, with 268 ROIs, it is more practical to summarize the results with the mean, standard error and 90% and 95% quantile intervals at each ROI. To demonstrate the results, here we illustrate the four main effects at the 268 parcels in the brain (Fig. 3): overall ISC, SRS, Sex, and Age. These effects can be interpreted in light of what is known from previous naturalistic scanning studies and the demographic and behavioral covariates of interest.

**Figure 3:**
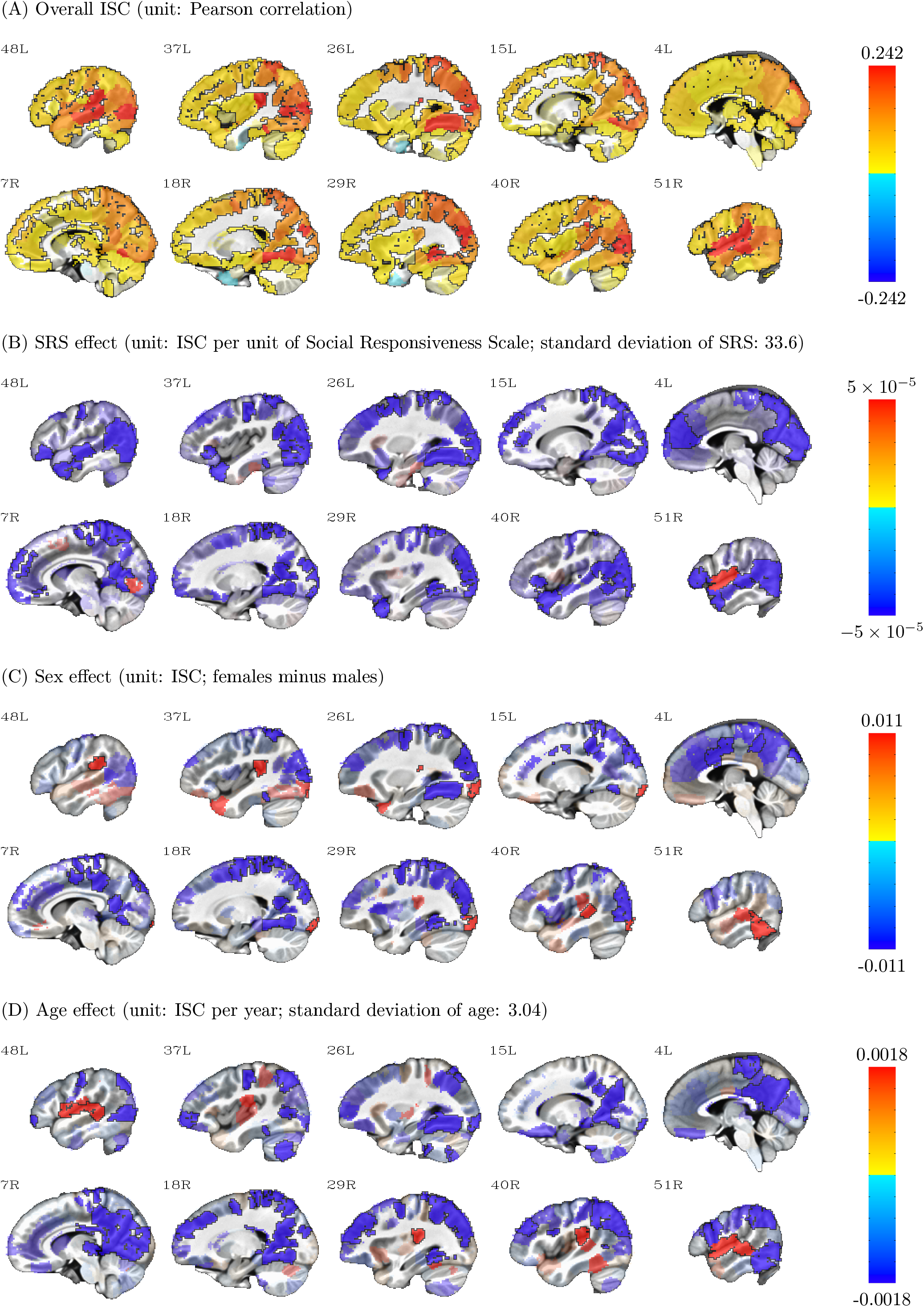
Four effects (overall ISC, SRS, Sex, and Age) derived from BML are shown here for the 268 parcels in sagittal view with slice numbers indicating the relative left-right location. Warm (or cold) colors show positive (or negative) effects, with the colorbar range set to the 95% quantile of the respective effect; effect opacity is determined by the posterior density: opaque regions are beyond 90% quantile tail (strong evidence), with transparency increasing toward the median (weak evidence). Note that the sex effect is shown as females minus males, meaning that in panel (C), blue regions show higher ISC in males while red regions show higher ISC in females.

First, much of the brain shows a substantial overall ISC effect (Fig. 3A). While this effect is particularly strong in primary visual and auditory cortex, there is evidence for synchrony in higher-order regions of association cortex as well. This is consistent with a large body of literature using naturalistic scanning to show that by exposing subjects to the same time-locked, complex, engaging stimulus, much of the brain becomes synchronized across subjects (Hasson et al., 2010).

Atop this general synchrony, our method revealed that subject-level covariates of interest affect the strength of ISC. In the case of Social Responsiveness Scale (SRS), most of these effects are negative (Fig. 3B), meaning that ISC is relatively stronger among children with low SRS scores than those with higher SRS scores. This is the expected direction given that lower SRS scores reflect better social function; in other words, children with good social skills are more synchronized while viewing a socially and emotionally evocative film as compared to children with more autistic traits and tendencies, corroborating previous reports (Hasson et al., 2009; Salmi et al., 2013; Byrge et al., 2015). There was substantial evidence for an effect in this direction in anterior and posterior regions along the midline as well as in temporal cortex, many of which are known to be involved in processing social information.

In the case of Sex (Fig. 3C), we observed higher ISC among males as compared to females in many posterior and central midline regions, as well as some visual association areas. In contrast, we observed higher ISC among females in the temporo-parietal junction and an inferior temporal region partially encompassing the fusiform gyrus.

In the case of Age (Fig. 3D), we observed that ISC generally declines with age, such that many regions (especially those in posterior midline and visual association regions) are more synchronized in younger children relative to older ones. One possible explanation for this is that idiosyncratic (i.e., subject-specific) responses emerge with age, leading to an increase in variance (and decrease in cross-subject synchrony) as children get older. Another potential explanation of these effects might be the choice of stimulus itself: the animated film may have been more engaging for younger subjects than older ones, who require more sophisticated content to fully capture their attention; future studies should explore the effect of stimulus on ISC values through development. The exception was a handful of regions along the superior temporal lobe, in which ISC increased with age. This may in part reflect language processes that are developed and refined as children mature, leading to more consistent responses among older subjects in these areas.

Beyond main effects, the BML framework also allows us to examine interaction effects among the covariates. We include one of these interactions, the Sex:Age interaction (Fig. 4), plus separate maps for the Age effect in each sex to facilitate interpretation (Fig. 4B, C). For example, a region in the inferior temporal lobe encompassing the fusiform gyrus seems to increase its ISC with age among females (Fig. 4C), while among males there is almost no evidence for such an age effect (Fig. 4B). Additionally, in some of the regions along the superior temporal lobe and insula, the increase in ISC with Age seems to be driven largely by females, which may reflect differing developmental trajectories in language and affect between the sexes.

**Figure 4:**
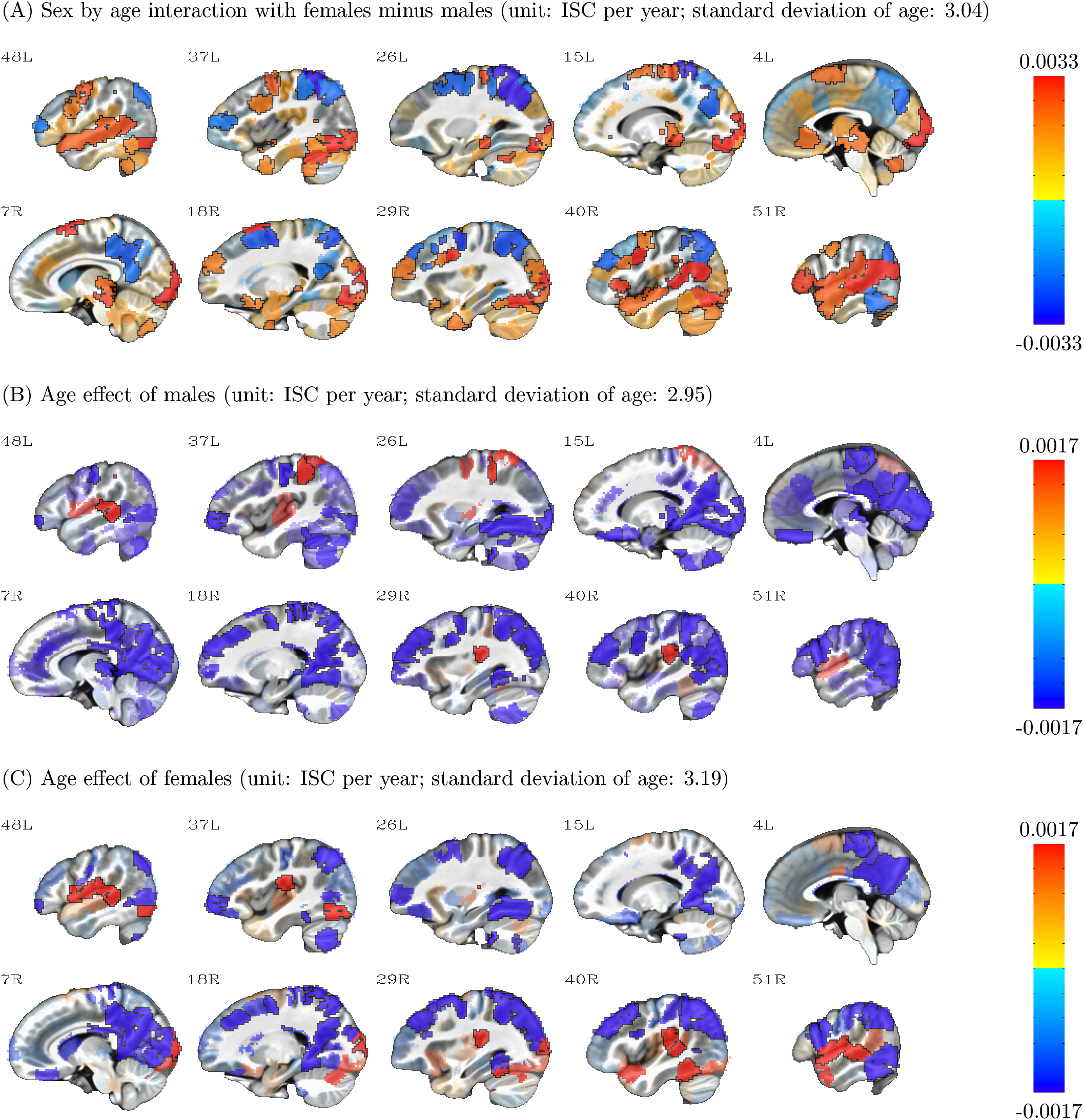
Interaction effects between sex and age derived from BML are shown here for the 268 parcels in sagittal view with slice numbers indicating the relative left-right location. Warm (or cold) colors show positive (or negative) effects, with the colorbar range set to the 95% quantile of the respective effect; effect opacity is determined by the posterior density: opaque regions are beyond 90% quantile tail (strong evidence), with transparency increasing toward the median (weak evidence). Note that the sex effect is shown as females minus males, meaning that in panel (1), blue regions show higher age effect in males while red regions show higher age effect in females.

One aspect in which ROI-based BML excels is the completeness and transparency in results reporting: if the number of ROIs is not overwhelming (e.g., less than 100), the summarized results for every ROI can be completely presented in a tabular form or in full distributions of posterior density (Chen et al., 2019). It is worth emphasizing that Bayesian inferences focus less on the point estimate of an effect and its associated quantile interval, but more on the whole posterior density that offers more detailed information about the effect uncertainty. Unlike the whole brain analysis in which the results are typically reported as the tips of icebergs above the water, posterior density reveals the extent of uncertainty regardless of strength of statistical evidence. In addition, one does not have to stick to a single harsh thresholding when deciding a criterion on the ROIs for discussion; for instance, even if an ROI lies outside of, but close to, the 95% quantile interval, it can still be reported and discussed as long as all the details are revealed. Such flexibility and transparency, as illustrated in Figures 3 and 4, are difficult to navigate or maneuver through the conventional cluster-based thresholding at the whole brain level.

## Discussion

Here, we introduce an extension to the LME platform, namely Bayesian multilevel modeling (BML), for jointly estimating inter-subject correlation during naturalistic scanning in a series of predefined regions. The advantages of this BML approach over previous approaches include: dissolution of multiplicity, ability to incorporate covariates, modeling efficiency, spatial specificity in outcome interpretation, results reporting and visualization.

We previously showed that the LME platform is relatively flexible for ISC data analysis at the whole brain voxel-wise level (Chen et al., 2017), compared to various nonparametric methods. Due to the complicated correlation structure among ISC values as illustrated in Fig. 1, ISC data analysis has been typically performed through a permutation approach of randomizing the temporal structure of EPI time series at the individual subject level as a null distribution for group analysis. Such an approach has been shown incapable of properly controlling for FPR. To improve the FPR controllability, subject-wise (SW) resampling has been developed (Chen et al., 2016) so that exchangeability and independence assumptions are satisfied, and the patterned correlation structure among the ISC values can be more accurately captured. Specifically, subject-wise bootstrapping (SWB) is a reasonable choice when inferring ISC for one group, while subject-wise permutation (SWP) testing is suitable for handling the comparison between two groups. Furthermore, to overcome the incapability of nonparametric approaches in incorporating explanatory variables, LME modeling with a crossed random-effects structure has been successfully employed (Chen et al., 2017) with each ISC value decomposed into fixed effects, two random intercepts that are associated with the two involved subjects, and residuals. Through a data duplication step that includes both the lower and upper triangular components of the ISC matrix as input, the LME approach achieves proper FPR controllability. In addition to the flexibility of incorporating subject-level explanatory variables, the LME framework allows for strong interpretation power, relatively low computational cost, and model quality control.

### ROI-based ISC analysis through BML as an extension of LME

The advantage of multilevel modeling lies in its capability of stratifying the data in a hierarchical or multilevel layout so that complex dependency or correlation structures can be properly accounted for coherently within a single modeling platform. Specifically applicable in the ISC context is a crossed or factorial layout across three crisscross layers, two sets of subject pairs and the list of ROIs. Even though the LME approach can quantitatively characterize the ISC effect of each subject pair as the combined effect of the respective subjects, the decomposition remains coarse. For instance, an LME model can accommodate neither the uniqueness of each subject pair nor that of each subject-ROI interaction, due to the LME system being potentially underdetermined from the overwhelming number of parameters. These limitations evince one motivation for our current work with BML as an extension to our previous work of LME modeling for ISC data analysis. That is, the idiosyncratic effect of each subject pair as well as that of each subject-ROI interaction can be modeled under BML since non-identifiability would be dissolved under BML because a Bayesian model can be identified as long as the posterior distribution is proper.

Applying the general BML modeling strategy (Chen et al., 2018) to the ISC context, we formulate the BML data generation mechanism for each dataset on a set of ROIs by extending an LME framework. Our adoption of BML, as illustrated with the demonstrative data analysis, indicates that BML holds some promise for ROI-based ISC data analysis and offers the additional advantages over traditional voxel-wise approaches:

1. Two multiplicity issues with the voxel-wise whole brain ISC analysis form another background for our work here. As for typical neuroimaging whole brain analysis, ISC analysis through LME would still have to face the multiplicity issue in the sense that the same model is applied as many times as the number of voxels. Therefore, correction for FWE would still have to be executed as part of the model or as an extra step. The popular approach of leveraging between cluster size and statistical strength has been widely adopted to control the overall FPR, but the penalty is usually too severe as the information shared across brain regions is not effectively considered in modeling (Chen et al., 2019 a). Another difficulty with the whole brain analysis is the sidedness issue in statistical testing. For a symmetric statistical distribution, one-sided testing for one direction (e.g., positive) would be justified if prior information is available regarding the sign of the test for a particular brain region. When no prior information is available for all regions in the brain, one cannot simply perform two separate one-sided tests in place of one two-sided test, and such a double-sidedness practice, although popularly practiced in the neuroimaging, warrants a Bonferroni correction because the two directions are independent with respect to each other (and each one-sided test is more liberal than a two-sided test at the same statistical evidence level). However, simultaneously testing both tails in tandem for whole brain analysis without correction for sidedness is widely used without clear justification, and this forms a source of multiplicity issue that needs proper accounting. Instead of separately correcting for multiple testing, BML incorporates multiple testing as part of the model by assigning a prior Gaussian distribution among the ROIs. In doing so, multiple testing is handled under the scaffold of the multilevel data structure by conservatively shrinking the original effect toward the center with the reasonable assumption that the effects among brain regions are usually similar and largely center within a finite range. In other words, instead of leveraging cluster size or statistical strength, BML leverages the commonality among ROIs through effective regularization, simultaneously achieving meaningful spatial specificity and detection efficiency. Even though the conventional correction for FWE in neuroimaging data analysis is considered desirable in controlling overblown false positives, it is not necessarily efficient nor practically meaningful to fight the strawman of absolutely zero effect anywhere in the brain. More importantly, arbitrary thresholding, regardless of the extent of rigor, artificially dichotomize the data, resulting in an undesirable situation: reporting only the results that pass thresholding unavoidably ignores the ones that may not differ much from the former. In addition, BML offers a flexible approach to dealing with double sidedness at the ROI level. When prior information about the directionality of an effect is available on some, but not all, regions (e.g., from previous studies in the literature), with the massively univariate approach for the whole brain one may face the issue of performing two one-tailed *t*-tests at the same time in a blindfold fashion. In contrast, the ROI-based BML approach disentangles the complexity since the posterior inference for each ROI can be made separately.
2. No duplication for input data is needed under BML. To keep a balanced data structure and to maintain proper overall FPR controllability under LME, we have to duplicate the input data with both the lower and upper triangular components of the ISC correlation matrix due to the fact those two sets of subject effects are parameterized as two separate parameter sets. In contrast, input data duplication under BML is unnecessary thanks to an implementation technique similar to the multi-membership modeling strategy available in the *R* package *brms* (Bürkner, 2017), halving the input data and the number of parameters for subject effects under BML, as opposed to LME.
3. BML may achieve higher spatial specificity through efficient modeling. A statistically identified cluster through the conventional whole brain analysis is not necessarily anatomically or functionally meaningful. In other words, a statistically identified cluster is not always aligned well with a brain region for diverse reasons such as “bleeding” effect due to contiguity among regions, and suboptimal alignment to the template space, as well as spatial blurring. In fact, investigators usually tabulate the location of the “peak” (i.e., maximum effect magnitude or statistic value) voxel for a cluster even though the cluster may only partially cover an anatomical region or overlap multiple brain regions or subregions. In contrast, the regions are utilized as prior spatial information, and the statistical inference for each region under BML is assessed by its effect strength relative to its peers, not by its spatial extent, providing an alternative to the conventional whole brain analysis with more accurate spatial specificity.
4. Full results reporting is possible for all ROIs under BML. The conventional NHST focuses on the point estimate of an effect supported with statistical evidence in the form of a *p*-value. In the same vein, typically the results from the whole brain analysis are displayed with sharp-thresholded maps or tables that only show the surviving clusters with peak statistic- or *p*-values. In contrast, as the focus under the Bayesian framework is on the posterior distribution, not the point estimate, of an effect, the totality of BML results can be summarized as shown in Figures ??. Such totality pits against the backdrop in which the effect estimates are usually not reported in the whole brain analysis (Chen et al., 2017b). In other words, BML modeling at the ROI level directly allows the investigator to present the effect estimate. More importantly, BML substantiates the reporting advantage not only because of modeling at the ROI level, but also due to the fact that the uncertainty associated with each effect estimate can be demonstrated in a much richer fashion. To some extent, the ROI-based BML approach can alleviate the arbitrariness involved in the thresholding with the current FPR correction practices. Even though BML allows the investigator to present the whole results for all regions, for example, in a table format, we do recognize that the investigator may prefer to focus the discussion on some regions with strong posterior statistical evidence. In general, with all effects reported in totality, regardless of their statistical evidence, the decision of choosing which effects to discuss in a paper should be based on cost, benefit, and probabilities of all results (Gelman et al., 2014). Specifically for neuroimaging data analysis, the decision still does not have to be solely from the statistical evidence; instead, we suggest that the decision be hinged on the statistical evidence from the current data, combined with prior information from previous studies. For example, one may still choose the 95% quantile interval as an equivalent benchmark to the conventional *p*-value of 0.05 when reporting the BML results. However, those effects with, say, 90% quantile intervals can still be utilized with a careful and transparent description, which can be used as a reference for future studies to validate or refute; or, such effects can be reported if they have been shown in previous studies. Moreover, rather than a cherry-picking approach on reporting and discussing statistically significant clusters in whole brain analysis^4^, we recommend a principled approach in results reporting in which the ROI-based results be reported in totality with a summary as shown in Figures ?? and be discussed through transparency and soft, instead of sharp, thresholding. We believe that such a highlighting and soft thresholding strategy is more healthy and wastes less information for a study that goes through a strenuous pipeline of experimental design, data collection, and analysis.
5. Inferences at the individuals are possible. As BML partitions the effect at subject pair level as the summation of multiple additive effects including the two involving subjects, the effect from each individual subject can be teased apart, revealing the contribution at the subject level as shown in formula (12), even though the input data for ISC analysis are at subject-pair level. Such effects at individual subject level could be beneficial as auxiliary information in exploring, for example, outlying subjects or association with behavior data.

### Limitations of ROI-based BML and future directions

ROIs can be defined through several ways depending on the specific study or information available regarding the relevant regions. For example, one can find potential regions involved in a task or condition including resting state and naturalistic scanning from the literature. Such regions are typically reported as the coordinates of a “peak” voxel (usually highest statistic value within a cluster), from which each region could be defined by centering a ball with a radius of, e.g., 6 mm in the brain volume (or by projecting an area on the surface). Regions can also be located through (typically coordinate-based) meta analysis with databases such as NeuroSynth (http://www.neurosynth.org) and BrainMap (http://www.brainmap.org), with tools such as brain_matrix (https://github.com/fredcallaway/brain_matrix), GingerALE (http://brainmap.org/ale), Sleuth (http://brainmap.org/sleuth), and Scribe (http://www.brainmap.org/scribe) that are associated with the database BrainMap. Anatomical atlases (e.g., http://surfer.nmr.mgh.harvard.edu, http://www.med.harvard.edu/aanlib) and functional parcellations (e.g., Shen et al., 2013; Schaefer et al., 2017) are some alternatives of region definition.

The limitations of the ROI-based BML are as follows.

1. Just as the FWE correction on the massively univariate modeling results is sensitive to the size of the full domain in which it is levied (whole brain, gray matter, or a user-defined volume), so the results from BML will depend to some extent on the number of ROIs (or which) ones included. For a specific ROI *j*, changing the composition among the rest of ROIs (e.g., adding an extra ROI or replacing one ROI with another) may result in different prior distributions and different posterior distributions even though most of the time the differences might be negligible. However, it merits noting that the regions should not be arbitrarily chosen but rather selected from the current knowledge and relevancy of the involving effect under investigation. More importantly, the impact of different number of regions on BML modeling is relatively small due to the adaptivity of the prior Gaussian distribution whose role is only for the general shape, not specific properties (i.e., mean and variance).
2. ROI data extraction involves averaging among voxels within the region. Averaging, as a spatial smoothing or low-pass filtering process, condenses, reduces or dilutes the information among the voxels within the region to one number, and loses any finer spatial structure within the ROI. In addition, the variability of extracted values across subjects and across ROIs could be different from the variability at the voxel level.
3. ROI-based analysis is conditional on the availability and quality of the ROI definition. One challenge facing ROI definition is the inconsistency in the literature due to inaccuracies across different coordinate/template systems and publication bias. In addition, some extent of arbitrariness is embedded in ROI definition; for example, a uniform adoption of a fixed radius may not work well due to the heterogeneity of brain region sizes. When not all regions or subregions currently can be accurately defined, or when no prior information is available to choose a region in the first place, the ROI-based approach may miss any potential regions if they are not included in the model.
4. The exchangeability requirement of BML assumes that no differential information is available across the ROIs in the model. Under some circumstances, ROIs can be expected to share differential information among some subgroups, especially when they are anatomically contiguous or more functionally related than the other ROIs (e.g., homologous regions in opposite hemisphere). Ignoring such hierarchical structure in the data, if substantially present, may lead to underestimated variability and inflated inferences. In the future we will explore the possibility of accounting for such a hierarchical correlation structure.

### Conclusion

Inter-subject correlation (ISC) captures the extent of the simultaneous synchronization at a brain region among a group of subjects who experience the same naturalistic setting such as movie watching or music listening. Extending our previous work of linear mixed-effects (LME) modeling, we adopt here an ROI-base Bayesian multilevel (BML) approach to decomposing each ISC effect into multiple additive effects. In addition to dissolving the multiplicity issue and achieving higher inference efficiency, the BML approach allows for full results reporting that pales in comparison with the prevalent adoption of dichotomous decision making under NHST, increasing transparency and reproducibility.

## Acknowledgments

The research and writing of the paper were supported (GC, PAT, PJM, PAB, RWC, ESF) by the NIMH and NINDS Intramural Research Programs (ZICMH002888) of the National Institutes of Health/HHS, USA. ESF is additionally supported by NIH grant K99MH120257. We thank the Child Mind Institute Healthy Brain Network for providing the data used here.

## Footnotes

1 The phenomenon is due to the following fact: with 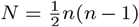 pairs of indices, there are totally 2*N* = *n*(*n* – 1) indices. When *n* is odd, each index repeats *n* – 1 times, thus they can be evenly distributed between the two sets after rearrangement because *n* – 1 is even; in contrast, when *n* is even, balance cannot be established because *n* – 1 is odd.

2 See https://en.wikipedia.org/wiki/Folded-t_and_half-t_distributions for the density *p*(*ν*, *μ*, *σ*^2^) of folded non-standardized *t*-distribution, where the parameters *ν*, *μ*, and *σ*^2^ are the degrees of freedom, mean, and variance.

3 The LKJ prior (Lewandowski, Kurowicka, and Joe, 2009) is a distribution over symmetric positive-definite matrices with the diagonals of 1s.

4 A popular cluster reporting method among the neuroimaging software packages is to simply present the investigator only with the icebergs above the water, the surviving clusters, reinforcing the illusionary either-or dichotomy under NHST.

## References

Abrams, D.A., Ryali, S., Chen, T., Chordia, P., Khouzam, A., Levitin, D.J., Menon, V., 2013. Inter-subject synchronization of brain responses during natural music listening. The European Journal of Neuroscience, 37(9):1458–69.

Alexander, Lindsay M., Jasmine Escalera, Lei Ai, Charissa Andreotti, Karina Febre, Alexander Mangone, Natan Vega-Potler, Nicolas Langer, Alexis Alexander, Meagan Kovacs, Shannon Litke, Bridget O’Hagan, Jennifer Andersen, Batya Bronstein, Anastasia Bui, Marijayne Bushey, Henry Butler, Victoria Castagna, Nicolas Camacho, Elisha Chan, Danielle Citera, Jon Clucas, Samantha Cohen, Sarah Dufek, Megan Eaves, Brian Fradera, Judith Gardner, Natalie Grant-Villegas, Gabriella Green, Camille Gregory, Emily Hart, Shana Harris, Megan Horton, Danielle Kahn, Katherine Kabotyanski, Bernard Karmel, Simon P. Kelly, Kayla Kleinman, Bonhwang Koo, Eliza Kramer, Elizabeth Lennon, Catherine Lord, Ginny Mantello, Amy Margolis, Kathleen R. Merikangas, Judith Milham, Giuseppe Minniti, Rebecca Neuhaus, Alexandra Levine, Yael Osman, Lucas C. Parra, Ken R. Pugh, Amy Racanello, Anita Restrepo, Tian Saltzman, Batya Septimus, Russell Tobe, Rachel Waltz, Anna Williams, Anna Yeo, Francisco X. Castellanos, Arno Klein, Tomas Paus, Bennett L. Leventhal, R. Cameron Craddock, Harold S. Koplewicz, and Michael P. Milham. 2017. An open resource for transdiagnostic research in pediatric mental health and learning disorders. Scientific Data, 4: 170181.

Amrhein, V., Greenland, S., 2017. Remove, rather than redefine, statistical significance. Nature Hum. Behav. 1:0224.

Bartels, A., Zeki, S., 2004. The chronoarchitecture of the human brain - natural viewing conditions reveal a time-based anatomy of the brain. NeuroImage 22, 419–433.

Browne, W.J., Goldstein H., Rasbash, J., 2001. Multiple membership multiple classification (MMMC) models. Statistical Modelling:103–124.

Bürkner, P., 2018. Advanced Bayesian Multilevel Modeling with the R Package brms.

Byrge, L., Dubois, J., Tyszka, J.M., Adolphs, R. and Kennedy, D.P., 2015. Idiosyncratic brain activation patterns are associated with poor social comprehension in autism. Journal of Neuroscience, 35(14), pp.5837–5850.

Cantlon, J.F., Li, R., 2013. Neural activity during natural viewing of Sesame Street statistically predicts test scores in early childhood. PLoS Biol. 11(1):e1001462. doi: 10.1371/journal.pbio.1001462.

Chen, G., Saad, Z.S., Britton, J.C., Pine, D.S., Cox, R.W., 2013. Linear Mixed-Effects Modeling Approach to FMRI Group Analysis. NeuroImage 73:176–190.

Chen, G., Shin, Y.-W., Taylor, P.A., Glen, D., Reynolds, R.C., Israel, R.B., Cox, R.W., 2016. Untangling the Relatedness among Correlations, Part I: Nonparametric Approaches to Inter-Subject Correlation Analysis at the Group Level. NeuroImage 142:248–259.

Chen, G., Taylor, P.A., Shin, Y.W., Reynolds, R.C., Cox, R.W., 2017. Untangling the Relatedness among Correlations, Part II: Inter-Subject Correlation Group Analysis through Linear Mixed-Effects Modeling. Neuroimage 147:825–840.

Chen, G., Xiao, Y., Taylor, P.A., Riggins, T., Geng, F., Redcay, E., 2019a. Handling Multiplicity in Neuroimaging through Bayesian Lenses with Multilevel Modeling. Neuroinformatics. doi: 10.1101/238998.

Chen, G., Bürkner, P.-C., Taylor, P.A., Li, Z., Yin, L., Glen, D.R., Kinnison, J., Cox, R.W., Pessoa, L., 2019b. An Integrative Approach to Matrix-Based Analyses in Neuroimaging. Human Brain Mapping (in press). doi: https://doi.org/10.1101/4595

Constantino, J.N., Gruber, C.P., 2012. Social responsiveness scale (SRS) (Western Psychological Services Torrance, CA).

Cox, D.R., Spjøtvoll, E., Johansen, S., van Zwet, W.R., Bithell, J.F., Barndorff-Nielsen, O., Keuls, M., 1977. The Role of Significance Tests. Scandinavian Journal of Statistics 4(2):49–70.

Cox, R.W. 1996. AFNI: software for analysis and visualization of functional magnetic resonance neuroimages, Computers and Biomedical Research, 29: 162–73.

Finn, E., Corlett, P., Chen, G., Bandettini, P., Constable, R., 2018. Trait paranoia shapes inter-subject synchrony in brain activity during an ambiguous social narrative. Nature Communications 9(1): 2043.

Fischl, B., 2012. FreeSurfer. Neuroimage, 62: 774–81.

Hasson, U., Nir Y., Levy, I., Fuhrmann, G., Malach, R., 2004. Intersubject synchronization of cortical activity during natural vision. Science 303:1634–1640.

Hasson, U., Yang, E., Vallines, I., Heeger, D.J., Rubin, N., 2008a. A hierarchy of temporal receptive windows in human cortex. Journal of Neuroscience 28(10):2539–2550.

Hasson, U., Landesman, O., Knappmeyer, B., Vallines, I., Rubin, N., Heeger, D.J., 2008b. Neurocinematics: The Neuroscience of Film. Projections Vol 2(1), 1–26.

Hasson, U., Avidan, G., Gelbard, H., Vallines, I., Harel, M., Minshew, N. and Behrmann, M., 2009. Shared and idiosyncratic cortical activation patterns in autism revealed under continuous realâĂŘlife viewing conditions. Autism Research, 2(4), pp.220–231.

Hasson, U., Malach, R., Heeger, D.J., 2010. Reliability of cortical activity during natural stimulation. Trends Cogn Sci, 14(1), 40–48.

Honey, C.J., Thomson, C.R., Lerner, Y., Hasson, U. (2012) Not lost in translation: Neural responses shared across languages. Journal of Neuroscience 32(44):15277–15283.

Kauppi, J.-P., Jääskeläinen, I.P., Sams, M., Tohka, J., 2010. Inter-Subject Correlation of Brain Hemodynamic Responses During Watching a Movie: Localization in Space and Frequency. Front Neuroinformatics 4: 5.

Kauppi, J.-P., Pajula J., Tohka, J., 2014. A Versatile Software Package for Inter-subject Correlation Based Analyses of fMRI. Frontiers in Neuroinformatics 8:2.

McShane, B.B., Gal, D., Gelman, A., Robert, C., Tackett, J.L., 2017. Abandon Statistical Significance. arXiv:1709.07588

Moraczewski, D., Chen, G., Redcay, E., 2018. Inter-subject synchrony as an index of functional specialization in early childhood. Scientific Reports 8(1) doi:10.1038/s41598-018-20600-0

Salmi, J., Roine, U., Glerean, E., Lahnakoski, J., Nieminen-von Wendt, T., Tani, P., LeppÃd’mÃd’ki, S., Nummenmaa, L., JÃd’Ãd’skelÃd’inen, I.P., Carlson, S. and Rintahaka, P., 2013. The brains of high functioning autistic individuals do not synchronize with those of others. NeuroImage: Clinical, 3, pp. 489–497.

Schmälzle, R., Häcker, F., Renner, B., Honey, C., Schupp, H., 2013. Neural correlates of risk perception during real-life risk communication. J. Neurosci. 33, 10340–10347.

Schmälzle, R., Häcker, F., Honey, C., Schupp, H., Hasson, U., 2015. Engaged listeners: shared neural processing of powerful political speeches. Soc Cogn Affect Neurosci 10(8): 1137–1143.

Shen, X., Tokoglu, F., Papademetris, X., Constable, R.T., 2013. Groupwise whole-brain parcellation from resting-state fMRI data for network node identification, Neuroimage 82: 403–15.

Smith, S.M., Nichols, T.E., 2009. Threshold-free cluster enhancement: addressing problems of smoothing, threshold dependence and localisation in cluster inference. Neuroimage. 44(1):83–98.

Wilson, S.M., Molnar-Szakacs, I., Iacoboni, M., 2008. Beyond Superior Temporal Cortex: Intersubject Correlations in Narrative Speech Comprehension. Cereb. Cortex 18(1): 230–242.

Yin, L., Xum X., Chen, G., Mehta, N.D., Haroon, E., Miller, A.H., Li, Z., Felger, J.C., 2019. Inflammation and decreased functional connectivity in depression: is there a ventral nexus? Under review.

Zhang, Y., Chen, G., Wen, H., Lu, K.-H., Liu, Z., 2017. Musical Imagery Involves Wernicke?s Area in Bilateral and Anti-Correlated Network Interactions in Musicians. Scientific Reports 7. doi:10.1038/s41598-017-17178-4

